# Susceptibility of Human Neural Stem Cells to SARS-CoV-2: Entry Mechanisms and Glycocalyx Influence

**DOI:** 10.1101/2025.11.10.687698

**Authors:** Ann Song, Cori Zuvia, Prue Talbot

## Abstract

Previous work demonstrated that ectodermal cells exhibit greater susceptibility to SARS-CoV-2 infection than endodermal and mesodermal cells, raising concerns about potential vulnerability of the developing nervous system. We hypothesized that neural stem cells (NSCs) derived from human ectoderm are susceptible to SARS-CoV-2 infection. Using pseudotyped viral particles representing both wild-type and Omicron spike variants, we confirmed efficient infection of NSCs, with Omicron variants preferentially utilizing endocytosis-mediated entry. Inhibition of endocytosis with filipin and OcTMAB significantly reduced infection across all spike variants. Low glycosylation levels on NSCs facilitated viral entry, and enzymatic removal of glycosylation increased their susceptibility. Ectodermal infection by SARS-CoV-2 raises serious concern for potential teratogenic effects on the nervous system, possibly causing latent or subclinical anomalies not immediately evident at birth. Therefore, future clinical studies and long-term surveillance of infants and children exposed *in utero* are necessary to investigate and identify potential neurodevelopmental deficits.

## Introduction

The human embryos and fetuses are highly susceptible to infectious agents such as viruses, which are well established teratogens (Adams Waldorf and McAdams, 2013). SARS-CoV-2 infection during early embryonic development is a significant concern, as embryos may not survive infection and those that survive may have severe developmental defects. Our previous research showed that human embryonic stem cells (hESCs), which model the epiblast (Nichols and Smith, 2009; 2011), and germ layer cells are susceptible to SARS-CoV-2 infection (Song et al. 2025). Susceptibility to infection varied among the different cell types with ectoderm being the most susceptible (Song et al. 2025). High levels of infection in ectodermal cells may lead to disturbances in neurodevelopment that may not be immediately apparent at birth.

In adults, neurological effects of SARS-CoV-2 are well documented and involve both the central and peripheral nervous systems, which originate from the ectoderm (Guerrero et al., 2021). Reported symptoms include anosmia and ageusia (Saniasiaya et al., 2020; Agyeman et al., 2020; Spudich and Nath, 2022), Guillain-Barré syndrome (Toscano et al., 2020; Rahimi, 2020), and encephalopathy (Poyiadji et al., 2020; Siahaan et al., 2022). Additionally, persistent neurological symptoms such as cognitive impairment (“brain fog”), fatigue, and sleep disturbances have been observed in individuals with long COVID (Perlis et al., 2022; Alkodaymi et al., 2022). Given our finding that the ectoderm is the most susceptible germ layer to SARS-CoV-2 infection (Song et al., 2025) paired with the established effects of SARS-CoV-2 on adult neural function, the derivatives of the ectoderm may be susceptible to SARS-CoV-2 infection.

SARS-CoV-2 effect on developing neural tissues is further emphasized by evidence of vertical transmission, which suggests a potential risk to ectodermal derivatives in the developing fetus. Vertical transmission of SARS-CoV-2 during the late third trimester is well documented, as evidenced by postpartum fetal samples (Vivanti et al., 2020; Hosier et al., 2020). Although some studies suggest this to be a rare case (Peng et al., 2020; Bwire et al., 2021), these studies may not have captured the optimal timing for qPCR results (Li et al., 2024a) or may have chosen non-responsive endpoints. The most robust evidence of vertical transmission has been reported in a first-trimester stillbirth study (Valdespino-Vázquez et al., 2021), which detected the virus in placental and fetal tissues, indicating the organ’s susceptibility to infection. Another first-trimester study involving SARS-CoV-2 positive pregnant women (Fenizia et al., 2022) reported that 30% of embryos/fetuses and 20% of syncytiotrophoblasts tested positive, which clearly indicate viral transmission through the placental membrane. The lack of information on ectodermal derivative infection further emphasizes the urgency of our research. Our study addresses this knowledge gap by utilizing a disease-in-a-dish model to demonstrate SARS-CoV-2 infection in earlier pregnancy stages.

In this study, neural stem cell (NSC), a derivative of the ectoderm, infection was examined with wild-type and Omicron-spike pseudoparticles (Song et al., 2023a; Song et al., 2025). NSCs were used to assess viral entry mechanisms, identify small molecule inhibitors that prevent infection, and investigate factors that cause tropism (differences in susceptibility among different cell types). Our findings provide insights into the susceptibility of early neural progenitors to SARS-CoV-2 and offer potential therapeutic strategies for mitigating the risk of vertical transmission.

## Methods

### Tissue Culture and Reagents

H9 hESCs (WiCell, Madison, WI) were grown in complete mTeSR Plus culture medium (STEMCELL Technologies, Vancouver, Canada) as described in our previous work (Lin and Talbot, 2011; Song et al. 2023a; Song et al., 2025). HEK 293T cells and an ACE2-overexpressing cell line (HEK 293T-ACE2) (ATCC, Manassas, VA) were grown in DMEM with high glucose and 10% FBS (Gibco, Carlsbad, CA) (Song et al. 2023 a, b; Song et al. 2025).

The following small molecule inhibitors were used to target TMPRSS2: ambroxol (10 µM; TCI Chemicals, Portland, OR), aprotinin (10 µM; Tocris, San Diego, CA), Camostat (10 µM; Sigma-Aldrich, Burlington, MA), Nafamostat (10 µM; TCI Chemicals, Portland, OR). Endocytosis inhibitors that were used included: Dyngo4a (20 µM, Selleckchem, Houston, TX), Pitstop2 (20 µM; Abcam, Cambridge, MA), OcTMAB (10 µM; Abcam, Cambridge, MA), MiTMAB (10 µM; Abcam, Cambridge, MA), mβCD (20 µM; Sigma-Aldrich, Burlington, MA), Nystatin (20 µM; Sigma-Aldrich, Burlington, MA), and Filipin (20 µM; Sigma-Aldrich, Burlington, MA).

### Neural Stem Cell Differentiation

To simulate early neural development, neural stem cell differentiation was performed with H9 hESCs using the STEMdiff^TM^ SMADi neural induction kit (STEMCELL Technologies, Vancouver, Canada) according to the manufacturer’s protocol. Single hESCs were first obtained using the Gentle Cell Dissociation Reagent (STEMCELL Technologies, Vancouver, Canada). 2 × 10^6^ cells were plated on Matrigel-coated 6-well plates in mTeSR Plus medium (STEMCELL Technologies, Vancouver, Canada) with ROCK inhibitor (10 µM; Tocris, San Diego, CA). Every 24 hours a medium change with neural induction medium was performed for 7 days. On the 7^th^ day, 2 × 10^6^ cells were dissociated using Accutase (Innovative Cell Technologies, San Diego, CA) then passaged onto a Matrigel-coated 6-well plate. Three passages were performed then cells were transferred to the STEMdiff^TM^ neural progenitor medium (STEMCELL Technologies, Vancouver, Canada) and cultured according to the manufacturer’s protocol.

### Fluorescence Microscopy

NSCs were seeded in 8-well chamber slides (Ibidi; Gräfelfing, Germany) and were fixed at room temperature in 4% paraformaldehyde in PBS for 15 min, then washed with PBS. For ACE2 and TMPRSS2 labeling, cells were treated with 50 mM DTT and 6 M guanidine-HCl for 5 minutes prior to quenching with 100 mM iodoacetamide. To verify that ACE2 and TMPRSS2 were located on the cell surface, fixed cells were labeled with the antibody and fluorescence lectin conjugate (WGA-FITC 1:100; Vector Laboratories, Newton, CA, USA).

For labeling with other antibodies, NSCs were permeabilized with 0.1% Triton X-100 for 10 min, then blocked with 10% donkey serum. Primary antibodies were diluted in blocking buffer. The following primary antibodies were used: mouse anti-PAX6 (1:60; Developmental Studies Hybridoma Bank, Iowa City, IA), mouse anti-NESTIN (1:200; Santa Cruz Biotechnology, Dallas, TX), mouse anti-OCT4 (1:200; Santa Cruz Biotechnology, Dallas, TX), goat anti-ACE2 (1:200; R&D Systems, Minneapolis, MN), and mouse anti-TMPRSS2 (1:200; Santa Cruz Biotechnology, Dallas, TX). Following overnight incubation with primary antibody at 4 °C, the cells were washed three times with 0.2% Tween in PBS, then incubated in the dark with Alexa Fluor-conjugated fluorescent secondary antibodies (1:500; Invitrogen, Carlsbad, CA). After washing in PBS, the cells were mounted with diluted Vectashield and DAPI for nuclear staining. A Nikon Eclipse Ti inverted microscope was used to image cells at 40x. NIS Elements Software and ImageJ software were used to collect images and analyze the data.

### TMPRSS2 Activity Assay

The TMPRSS2 cleavage assay was performed as described in our previous work (Song et al. 2025). A TMPRSS2 fluorogenic substrate, Boc-Gln-Ala-Arg-AMC HCl (2.5 mM, Bachem, Torrance, CA), was prepared in 50 mM Tris (pH 8) and 150 mM NaCl. After differentiation, NSCs were washed twice with PBS and lysed for 1 min on ice in RIPA buffer. The cells were then sheared with a 21-gauge needle, followed by centrifugation at 3,000 rpm for 5 min at 4 °C. The lysate protein was quantified using the Pierce BCA assay kit (Thermo Scientific, Waltham, MA). 10 µg of protein was added to each reaction well. Cells were preincubated with TMPRSS2 inhibitors for 24 h before measuring enzymatic activity. The fluorogenic substrate was added to each well at a final concentration of 10 µM. Fluorescence intensity was measured at 340/440 nm using a BioTek Synergy HTX, multi-mode microplate reader (Winooski, VT).

### Lentiviral Production to Generate SARS-CoV-2 Pseudoparticles

SARS-CoV-2 pseudoparticles were generated as described in our previous work (Song et al., 2023a, b; Song et al. 2025). HEK293T cells were plated with antibiotic-free medium at a density of 7 × 10^6^ cells in a T75 flask and transfected using lentiviral plasmids (Table 1) and a Lipofectamine3000 Kit (Invitrogen, Carlsbad, CA) according to the manufacturer’s protocol. After overnight incubation, fresh medium was added to cells supplemented with 1% BSA. Conditioned medium was collected and centrifuged 48 h post-transfection. The supernatant was filtered using a 0.45 µm Acrodisc syringe filter (Cytivia Life Sciences, Marlborough, MA), and the filtrate was mixed with 5x polyethylene glycol (Abcam, Cambridge, UK) and precipitated overnight at 4°C. The lentivirus was collected by centrifugation, and the pellet was resuspended in Viral Re-suspension Solution (Abcam, Cambridge, UK). Virus aliquots were stored at −80°C. Prior to use in experiments, the transduction efficiency of each batch of viruses was tested in HEK 293T-ACE2.

**Table 1.** List of lentivectors and their amounts for a 75-cm^2^ flask.

| Plasmid Name | Amount (μg) |
| --- | --- |
| pHAGE2-CMV-ZsGreen-W | 7.5 |
| HDM-Hgpm2 | 1.65 |
| pRC-CMV Rev1b | 1.65 |
| HDM-tat1b | 1.65 |
| pcDNA3.3_CoV2_D18 (wild-type)<br>or<br>pcDNA3.3_SARS2_omicron_BA.1<br>(omicron BA.1 variant) or<br>pcDNA3.3_SARS2_omicron_BA.2<br>(omicron BA.2 variant) | 2.55 |

### Pseudotyping of human SARS-CoV-2 viral pseudoparticles

SARS-CoV-2 pseudoparticle infection was performed as described in our previous work (Song et al., 2023a, b; Song et al. 2025). 1.25 × 10^5^ of NSCs were seeded in 48-well plates, then infected 24 hours later with SARS-CoV-2 viral pseudoparticles using a multiplicity of infection (MOI) of 0.1. H9 hESCs were used as an undifferentiated control. The medium was replaced after overnight incubation of the infected cells. Cells were dissociated with Accutase 48 h post-infection and washed three times with 0.5% BSA in PBS. After the final wash, the cells were resuspended in the same wash buffer and analyzed using flow cytometry. A Novocyte flow cytometer (Agilent Technologies, Santa Clara, CA) was used to detect ZsGreen in the FITC channel. The resulting flow cytometer files were analyzed using NovoExpress software. Mock infection was used as a background control. Prior to infection, the cells were preincubated for 2 h with TMPRSS2 inhibitors or endocytosis inhibitors, which were kept in the medium until harvesting for flow cytometry.

### Endocytosis Assay

NSC differentiation was performed in 8-well chamber slides (Ibidi; Gräfelfing, Germany). The cells were incubated overnight with TRITC-conjugated dextran (Invitrogen, Carlsbad, CA) with and without endocytosis inhibitors. After fixation in 4% paraformaldehyde, mounting was performed using diluted Vectashield with DAPI, and images were collected using a Nikon Eclipse inverted microscope.

### Cell surface glycosylation and deglycosylation assay

To determine surface glycosylation patterns, NSCs were labeled with various lectins (Con A, DBA, PNA, RCA120, SBA, UEA I, WGA from Vector Laboratories, Newton, CA, USA). After lectin labeling, cells were fixed in 4% paraformaldehyde for 15 minutes at room temperature and mounted in Vectashield containing DAPI for microscopy.

For deglycosylation, NSCs were seeded in 48-well plates. After 24 hours, the cells were treated with neuraminidase (Sigma-Aldrich, Burlington, MA) in DMEM at 37°C for variable periods of time (0 min, 45 min, 90 min, 180 min, 360 min). 1 Unit of neuraminidase liberates 1.0 µM of N-acetyl neuraminic acid per minute at 37°C using bovine submaxillary mucin. After deglycosylation, neuraminidase was quenched with culture medium and cells were washed with PBS.

For infection, cells were transferred to neural maintenance medium, and SARS-CoV-2 pseudoparticles were added after the glycosidase treatment to allow infection. After 48 hours, cells were collected and analyzed for infection using flow cytometry.

### Data Analysis and Statistics

For infection and TMPRSS2 activity data, the mean and standard error of the mean for three independent experiments were plotted using GraphPad Prism 10 software (GraphPad, San Diego, CA). For infection data, the mean of the DMSO group was set to 100, and the inhibitor groups were compared to this value. Statistical significance was determined using Minitab Statistics Software (Minitab, State College, PA). When the data were not normally distributed, they were subjected to a logarithmic, arcsine square root, or Box-Cox transformation, and the data were retested to confirm that they satisfied the analysis of variance (ANOVA) model (normal distribution and homogeneity of variances). Infection analyses were performed using a one-way ANOVA, while TMPRSS2 cleavage analysis was performed using a two-way ANOVA, in which the factors were time and treatment. When the ANOVA means were significant (p < 0.05), groups treated with inhibitors were compared to the control group using Dunnett’s or Tukey post hoc analysis.

## Results

NSC differentiation from H9 hESCs was successful as the pluripotency marker OCT4 was absent and the neural markers PAX6 and NESTIN were present (Figures 1A-C). WGA, a cell surface-binding lectin (Figures 1D and H), and ACE2 and TMPRSS2, the machinery for SARS-CoV-2 entry, were present and often aggregated into clusters on NSCs (Figures 1E and 1I). Surface localization of ACE2 and TMPRSS2 was confirmed by colocalization with WGA, which generated yellow fluorescence in clustered regions (Figures 1F and 1J). ACE2 and TMPRSS2 were localized in the plasma membrane both as clusters above the nuclei and as individual proteins (small puncta) away from the nuclei (Figures 1G and 1K). In addition, ACE2 and TMPRSS2 were often colocalized as indicated by the yellow fluorescence in many of the puncta in the merged images (Figures 1 L-N).

**Figure 1.**
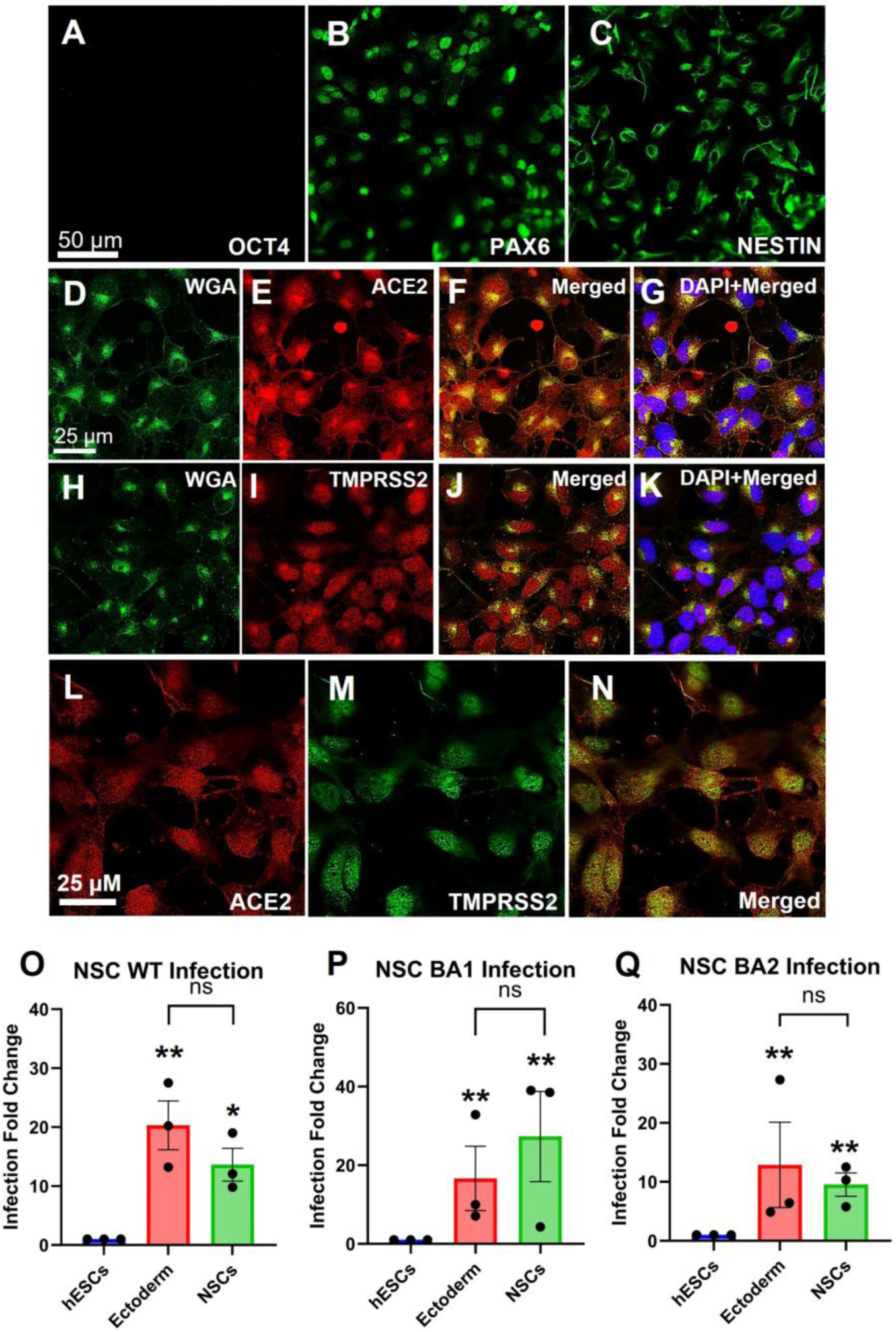
NSCs were susceptible to SARS-CoV-2 pseudoparticle infection. Immunocytochemistry showing successful differentiation of H9 hESCs into NSCs using (A) OCT4, (B) PAX6, and (C) NESTIN labeling. Immunocytochemistry showing (E) ACE2 and (I) TMPRSS2 localization in NSCs. Both samples were labeled with WGA conjugated to FITC to verify surface localization of (D) ACE2 or (H) TMPRSS2. Merged images of (F) WGA and ACE2 and (J) WGA and TMPRSS2 show colocalization. Colocalization of (L) ACE2 and (M) TMPRSS2 is shown in (N) the merged image. Representative images of three independent experiments are shown. Flow cytometry was performed to check the relative infection in NSCs with (O) WT Spike pseudoparticles, (P) BA1 pseudoparticles, and (Q) BA2 pseudoparticles. One-way ANOVAs were performed on raw data, and Tukey post-hoc test was used to compare to the hESCs control. Data are the means ± SEM of three independent experiments. * = p < 0.05, ** = p < 0.01.

For each spike protein pseudoparticle (WT, BA1, and BA2), the ectoderm was significantly infected (Figures 1O-Q), in agreement with our prior research on WT infection (Song et al. 2025) and further showing that the BA1 and BA2 variants also infected ectoderm. NSCs were also successfully infected by the three types of spike pseudoparticles, and their percentage of infection was significantly elevated compared to the hESC control (Figures 1O-Q). Infection of the ectoderm and NSCs did not differ significantly from each other for any of the variants.

Inhibition of TMPRSS2 cleavage activity was evaluated using small molecule inhibitors (camostat, nafamostat, aprotinin, and ambroxol) (Figure 2A). All inhibitors successfully decreased TMPRSS2 cleavage of the fluorogenic substrate with inhibition ranging from 40-50% (Figure 2A).

**Figure 2.**
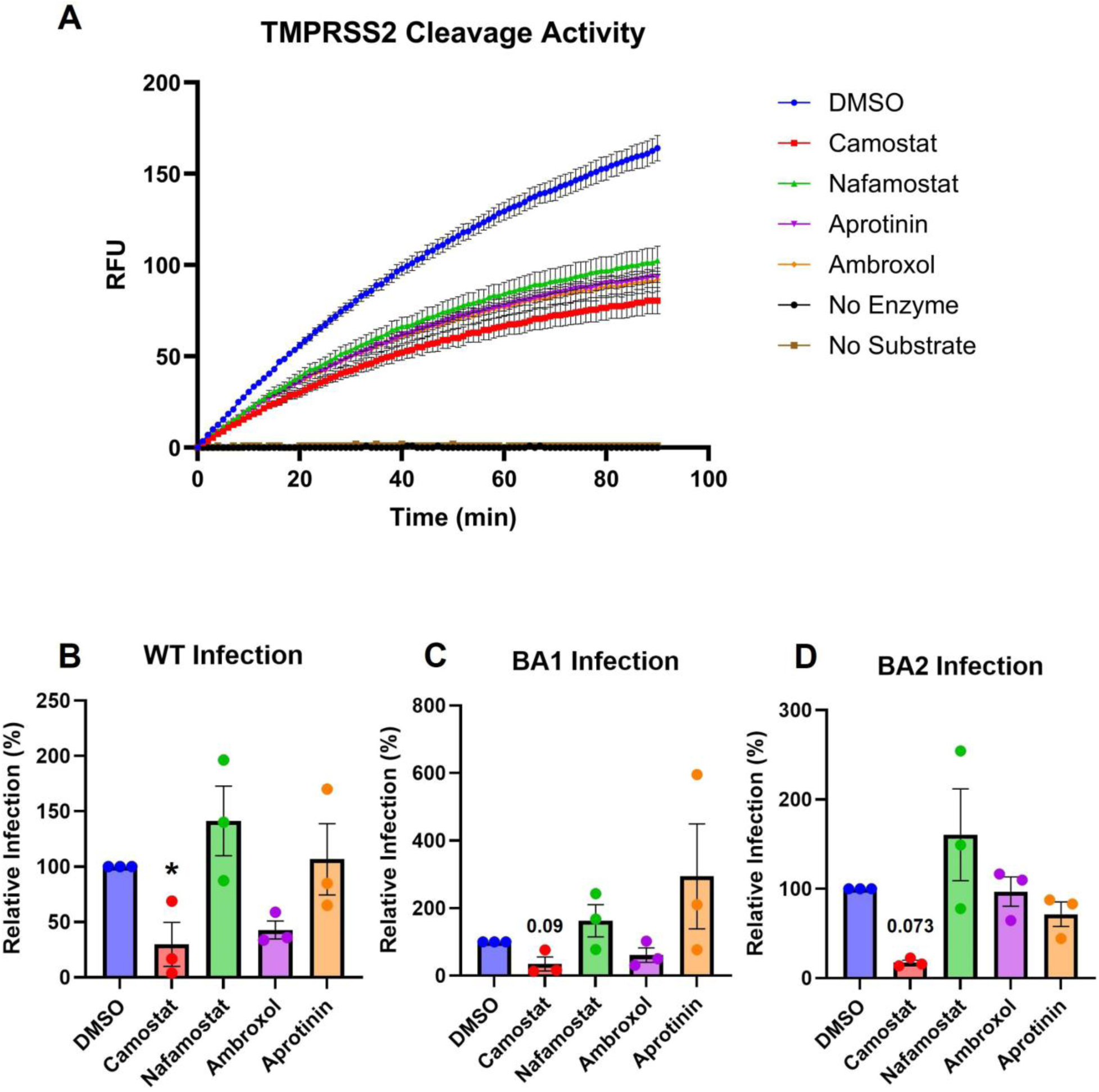
TMPRSS2 inhibitors decreased infection in NSCs. (A) TMPRSS2 activity of NSCs. RFU = relative fluorescence units. Camostat significantly decreased infection with (B) WT Spike pseudoparticles, (C) BA1 pseudoparticles, and (D) BA2 pseudoparticles. In (B-D), data are normalized to the DMSO control group. Each group is the mean ± SEM of three independent experiments. * = p < 0.05.

Infections were performed in the presence of TMPRSS2 inhibitors, and the data were graphed with the DMSO control set to 100% (Figures 2B-D). Camostat decreased infection in all three pseudoparticle groups; however, it only reached significance in the WT (Figures 2B-D). In some cases (e.g., nafamostat), inhibitors appeared to increase infection; however, these increases were not significantly different than the DMSO.

SARS-CoV-2 can also enter cells via endocytosis. Dextran conjugated with TRITC was used to visualize endocytosis in the presence or absence of inhibitors (Dyngo4A, OcTMAB, MiTMAB, Pitstop2, mβCD, nystatin, filipin) (Figure 3A). The untreated DMSO control successfully endocytosed dextran-TRITC, which appeared as small red puncta inside the cells. NSCs treated with endocytosis inhibitors showed decreased red fluorescence, indicating reduced endocytosis (Figure 3A). NSCs treated with Dyngo4A, OcTMAB, MiTMAB, Pitstop2, nystatin, and filipin showed little fluorescence, whereas mβCD showed some endocytic uptake, indicating less efficient inhibition (Figure 3A).

**Figure 3.**
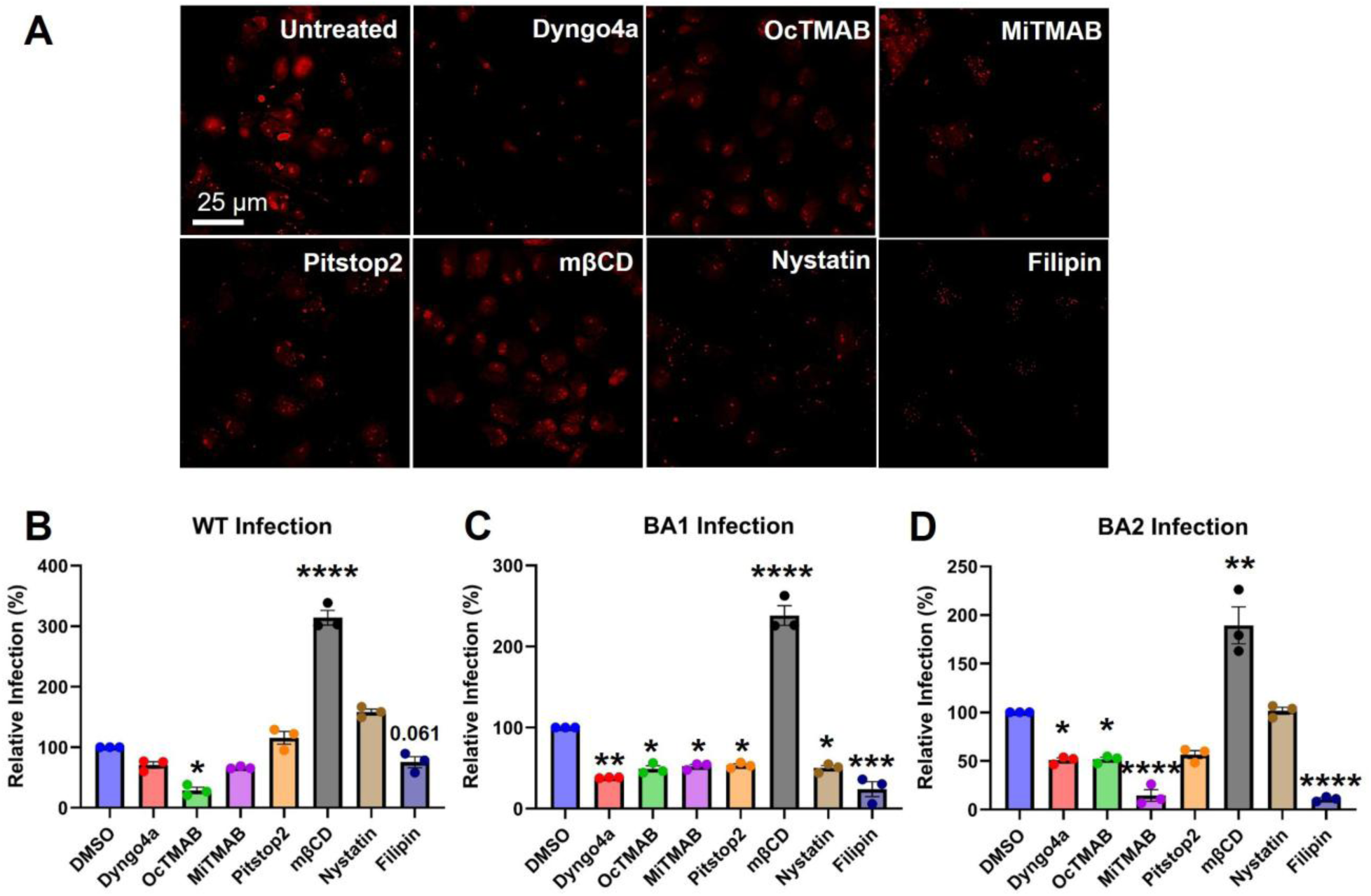
Endocytosis inhibitors decreased infection in NSCs. (A) All endocytosis inhibitors decreased endocytosis of TRITC conjugated dextran. (B) OcTMAB and Filipin significantly decreased infection with WT Spike pseudoparticles. (C) Dyngo4a and Filipin significantly decreased infection with BA1 pseudoparticles. (D) Dyngo4a, OcTMAB, MiTMAB, Pitstop2, and Filipin significantly decreased infection with BA2 pseudoparticles. In (B-D), one-way ANOVAs were performed on the raw infection data, and Dunnett’s post-hoc test was used to compare treated groups to the DMSO control. Data are the means ± SEM of three independent experiments. * = p < 0.05, ** = p < 0.01, *** = p < 0.001, **** = p < 0.0001.

Infection was next performed with the three types of pseudoparticles and all endocytosis inhibitors (Dyngo4a, OcTMAB, MiTMAB, Pitstop2, Filipin, mβCD, nystatin). Infection of the DMSO control was set to 100% and the inhibitor groups were compared to the control (Figure 3B-D). All endocytosis inhibitors (Dyngo4a, OcTMAB, MiTMAB, Pitstop2, and Filipin) decreased infection relative to the DMSO control, with the exceptions of mβCD and nystatin. For WT spike pseudoparticles, OcTMAB was significantly lower than the DMSO control and filipin was close to significance (p < 0.061) (Figure 3B). For BA1 spike pseudoparticles, Dyngo4a and filipin were significantly lower than the control (Figure 3C). For BA1 pseudoparticles, all inhibitors were significantly lower than the DMSO control except mβCD, which was higher (Figure 3C). For BA2 spike pseudoparticles, all inhibitors were significantly lower than the control except for mβCD and nystatin (Figure 3D). Filipin consistently and significantly reduced infection compared to the DMSO in WT, BA1, and BA2 (*p* = 0.061, *p* < 0.001, *p* < 0.0001), in agreement with the fluorescent assay in which filipin was the most effective drug at inhibiting dextran uptake.

Although the uptake of dextran with mβCD was reduced compared to that of the untreated control (Figure 3A), evidence of endocytosis was observed in the infection assay. mβCD consistently elevated the relative infection levels compared to DMSO and was significant for WT and BA2 (*p* < 0.05, *p* < 0.01), indicating that mβCD is not an effective inhibitor of SARS-CoV-2 viral entry via endocytosis. Although nystatin increased infection in WT and BA2, these means were not statistically significant. However, nystatin did significantly decrease BA1 infection.

To determine if the cell surface glycocalyx affects infectability of the NSCs, glycosylation was first assessed using FITC-conjugated lectins (Con A, RCA 120, WGA, UEA I, PNA, SBA, and DBA) (Figure 4A-G). Con A, RCA120, and WGA bound to NSCs (Figures 4A-C), whereas UEA I, PNA, SBA, and DBA showed little to no binding (Figure 4D-G). Based on our previous study (Song et al. 2025), we hypothesized that removing glycosylation would increase infectability by providing better access to the ACE2 receptor on the host cell plasma membrane. NSCs were incubated with 0.006 U/mL or 0.018 U/mL of neuraminidase for 0, 45, 90, 180, or 360 min to remove surface sialic acid. Neuraminidase treatment decreased WGA-FITC binding (Figure 4H), indicating successful removal of sialic acid groups from the cell surface.

**Figure 4.**
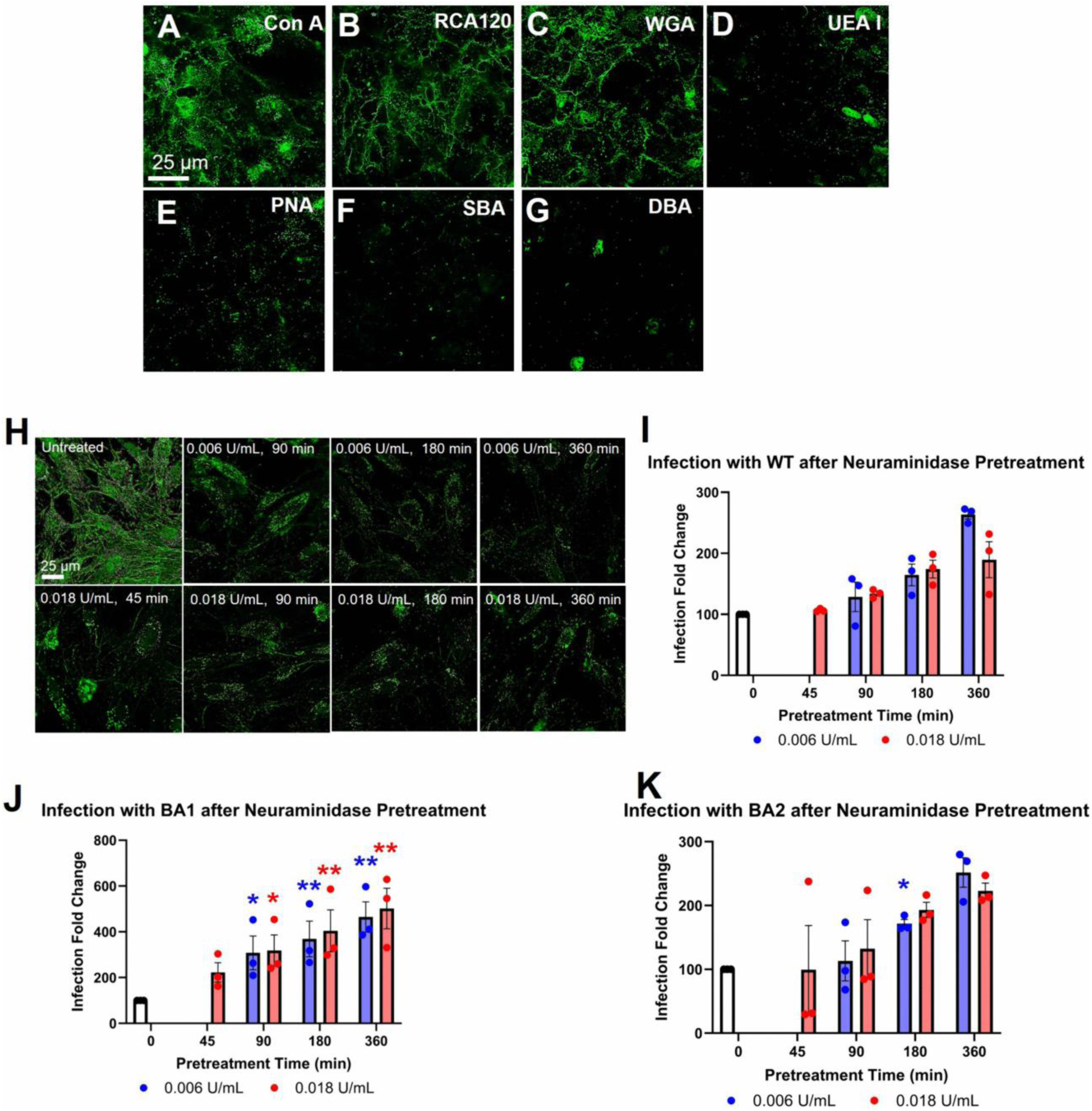
Neuraminidase treatment of NSCs increased infection. (A-G) Glycosylation pattern of NSCs was characterized by labeling with FITC-conjugated lectins (ConA, RCA120, WGA, UEA I, PNA, SBA, DBA). NSCs were also labeled with or without neuraminidase treatment (0.006 U/mL, 0.018 U/mL).

Treated cells showed loss of WGA-FITC labeling versus the untreated control. (H) Neuraminidase treatment significantly increased infection with: (I) WT, (J) BA1, and (K) BA2 pseudoparticles, based on one-way ANOVAs on raw infection data followed by Dunnett’s post-hoc tests to compare neuraminidase treated groups to the untreated control. Data are the means ± SEM of three independent experiments. * = p < 0.05, ** = p < 0.01, **** = p < 0.0001. Blue asterisk = one-way ANOVA significance for 0.006 U/mL of neuraminidase; Red asterisk = one-way ANOVA significance for 0.018 U/mL of neuraminidase.

To determine if removal of sialic acid would increase infection of hESCs, cells were pretreated with neuraminidase (0.006 U/mL, 0.018 U/mL) for various periods of time before infection with SARS-CoV-2 pseudoparticles. For WT and BA2 spike pseudoparticles, an overall increase in infection was observed in the 0.006 U/mL groups, and BA2 pretreated for 180 min was significantly increased; however, in the 360 min groups, infection was not significantly different from the control, probably because some cell death occurred (Figures 4 I, K). For BA1 spike pseudoparticles, infection significantly increased as the time of pretreatment with neuraminidase increased, with the higher concentration producing a larger effect faster (Figure 4J). Overall, a positive correlation between the loss of WGA labeling and an increase in infection was observed for the three types of pseudoparticles. Fold increases ranged from about 2-4 with BA2 pseudoparticles having the largest increase.

## Discussion

The teratogenic potential of SARS-CoV-2 remains unclear, as our current understanding is based predominantly on speculative interpretations of limited reports on vertical transmission (Youngster et al., 2022). Given that viral teratogens often target the nervous system with developmental effects sometimes manifesting later in life (Hernandez-Diaz et al., 2022), this gap in knowledge carries urgent implications. Our study addresses this concern by demonstrating the impact of SARS-CoV-2 on cells present in early neural development. We adapted our disease-in-a-dish model to investigate the infection of NSCs with wild type and Omicron spike (BA1, BA2) pseudoparticles (Song et al., 2023a; Song et al., 2025). We characterized the SARS-CoV-2 entry mechanisms and identified small molecule inhibitors that can prevent infection. Our findings demonstrate the susceptibility of NSCs to SARS-CoV-2 infection and suggest this virus is a potential teratogen. Notably, NSCs utilized both membrane fusion and endocytosis entry pathways, with Omicron pseudoparticles showing a marked preference for endocytosis. Because NSCs were more susceptible to SARS-CoV-2 infection than hESCs, we predicted that NSCs would have low levels of glycosylation, as seen previously with ectodermal cells (Song et al., 2025). Our results with hESCs, ectoderm, and NSCs show that cells with a reduced glycocalyx are significantly more susceptible to SARS-CoV-2 infection (Song et al., 2025). Partial removal of the glycocalyx on NSCs significantly increased infection with each type of pseudoparticle, supporting our previous data that showed heavy glycosylation on embryonic cells impeded infection (Song et al., 2025).

Our data provide strong support for the hypothesis that NSCs, like their ectodermal progenitors (Song et al., 2025), are highly susceptible to SARS-CoV-2 pseudoparticle infection. Building on our previous work, which linked ectodermal vulnerability to elevated TMPRSS2 activity, dual utilization of viral entry pathways, and a sparse glycocalyx, we now demonstrate that NSCs share these permissive features. The high infectability observed with NSCs further addresses viral tropism that likely stems from two principal factors: utilization of both viral entry pathways and the minimal glycocalyx on the NSCs surface. Together, these attributes create a cellular microenvironment favorable to viral invasion, raising the possibility of SARS-CoV-2-induced developmental perturbations in ectodermal derivatives, including the central nervous system and integumentary structures.

Adult derivatives of the ectoderm including central nervous system cells, such as NSCs, may have increased susceptibility to SARS-CoV-2. This idea is supported by clinical reports of anosmia and ageusia (Saniasiaya et al., 2020; Agyeman et al., 2020) in asymptomatic adult COVID-19 patients (Paderno et al., 2020; Niazkar et al., 2020), suggesting SARS-CoV-2 damage to the olfactory epithelium (Meunier et al., 2020). The medical literature also documents the involvement of the nervous system in COVID-19 (Guerrero et al., 2021) with SARS-CoV-2 virus detected in the brains of deceased patients (Puelles et al., 2020) and in cerebrospinal fluid during autopsies (Yavarpour-Bali and Ghasemi-Kasman, 2020). The expression of ACE2 in the brain, even at low levels, supports the neuroinvasive potential of SARS-CoV-2 (Li et al., 2020b; Baig et al., 2020). Neurological complications in adults should not be underestimated, as conditions such as Guillain-Barre syndrome (Poyiadji et al., 2020) and in rare instances, acute encephalopathy (Toscano et al., 2020) have apparently been triggered by SARS-CoV-2 infection. Taken together, these findings in conjunction with our data suggest that SARS-CoV-2 can target ectodermal derivatives across developmental stages. Accordingly, it is critical that clinicians monitor infants and children born to mothers infected with SARS-CoV-2 during pregnancy for signs of cognitive or neurodevelopmental impairment.

This need for neurological monitoring is underscored by persistent symptoms reported in individuals with post-acute sequelae of SARS-CoV-2 infection (“Long COVID”), including fatigue, cognitive dysfunction (“brain fog”), dyspnea, and sleep disturbances (Perlis et al., 2022; Alkodaymi et al., 2022). A recent study quantified cognitive impairment in affected individuals, revealing a measurable IQ loss in long-COVID patients experiencing brain fog (Hampshire et al., 2024). While overt anatomical deficits have not been reported in children born to COVID mothers infected with SARS-CoV-2 during pregnancy (Hernandez-Diaz et al., 2022), in contrast to the microcephaly observed in congenital Zika virus infection (Brasil et al., 2016), subtle alternations in brain function and behavior may nevertheless exist in children exposed to COVID *in utero*. However, a recent large electronic health record cohort corroborated clinical concerns by reporting that maternal SARS-CoV-2 infection during pregnancy was associated with increased risks of a neurodevelopmental diagnosis by age three (adjusted OR 1.29, 95% CI 1.05–1.57) (Shook et al., 2025). Emerging experimental and clinical evidence support the idea that neuroectodermal development is adversely affected by SARS-CoV-2, which is vertically transmitted from an infected mother to her embryo/fetus.

Omicron was rapidly established as a precursor to future dominant SAR-CoV-2 strains based on its heavy mutations, increased transmissibility, and its capacity to cause breakthrough infections in vaccinated populations (Wei et al., 2021; Cai et al., 2023; Mohanty et al., 2022; Manchanda et al., 2023; Ou et al., 2022). At the time of writing, the XEC subvariant of Omicron had emerged as the predominant circulating strain (Aljabali et al., 2025). This variant is distinguished by its enhanced immune evasion attributed to spike protein mutations that significantly diminish neutralizing antibody responses in vaccinated individuals (Chan et al., 2023; Shah and Woo, 2022; Hoffmann et al., 2022; Fossum et al., 2024; Li et al., 2024; Scarpa et al., 2025). Our results demonstrate that NSCs exhibit marked susceptibility to both BA1 and BA2 Omicron pseudoparticles, emphasizing concerns about viral tropism toward ectodermal derivatives. Given Omicron’s virulence and resistance to pre-existing immunity, heightened vigilance is warranted among pregnant individuals, as *in utero* exposure may pose risks to fetal neurodevelopment.

Our study investigated viral entry as a potential therapeutic target, consistent with prior proposals emphasizing host-cell entry modulation (Mercer et al., 2010; Mazzon and Marsh, 2019). Our data indicate that Omicron BA1 and BA2 pseudoparticles preferentially use endocytosis-based entry, in agreement with other studies that distinguish Omicron from earlier variants, such as Alpha, Beta, and Delta (Emanuel et al., 2023; Pennisi et al., 2025; Iwata-Yoshikawa et al., 2022). This shift in entry preference has been attributed to spike protein mutations that enhance endocytic uptake while diminishing TMPRSS2-dependent membrane fusion (Yamamoto et al., 2022). These mechanistic insights suggest that therapeutic approaches targeting the endocytosis pathway may offer improved efficacy against Omicron infection (Song et al., 2023b; 2025).

Our study demonstrated that both OcTMAB and filipin consistently inhibited NSC infection across all cell types of SARS-CoV-2 pseudoparticles. OcTMAB, a type of quaternary ammonium salt predicted to be a SARS-CoV-2 inhibitor (Hessien et al., 2023) by interference with dynamin-dependent endocytosis (Quan et al., 2007), is widely used in disinfectants recommended for viral decontamination, supporting its antiviral properties (Hora et al., 2020). A materials engineering study found that polypropylene composites containing an OcTMAB analog (CTAB) have antiviral SARS-CoV-2 activity, suggesting utility in the development of antiviral materials such as bags or containers (Guerrero-Bermea et al, 2022). Filipin similarly limited viral entry by disrupting caveolae-mediated endocytosis, a pathway reliant on cholesterol-rich membrane domains essential for SARS-CoV-2 internalization (Song et al., 2023b; Alkafaas et al., 2022). Consistent with our findings, Zhou et al. (2022) reported that filipin blocked SARS-CoV-2 entry into Vero E6 cells. The significant reduction in viral entry by filipin treatment highlights its potential as a targeted antiviral agent acting through endocytic pathways (Song et al., 2023b; Pirzada et al., 2021). The strong inhibitory effects of OcTMAB and filipin on endocytosis demand further investigation to establish their translational potential for preventing SARS-CoV-2 infection of the nervous system.

While OcTMAB and filipin effectively reduced SARS-CoV-2 pseudoparticle infection in NSCs, not all inhibitors of caveolae-mediated endocytosis yielded protective effects. Notably, mβCD not only failed to inhibit endocytosis but paradoxically enhanced infection, highlighting the need for caution in the application of endocytosis inhibitors. Our data are consistent with prior studies in which mβCD did not inhibit pathogen uptake by endocytosis. For instance, mβCD treatment did not prevent herpes simplex virus type 1 entry into human keratinocytes (Rahn et al., 2011) or Campylobacter jejuni internalization in HeLa cells (Konkel et al., 2013). mβCD also has non-specific effects on cellular homeostasis, which in severe cases, can lead to neuronal death (Dai et al., 2017) or disrupt synaptic transmission and plasticity by altering membrane lipid organization and affecting receptor function (Korinek et al., 2015).

Our previous study showed that the glycocalyx modulates the infectability of human embryonic cells (Song et al., 2025). In the current study, glycosylation profiling further demonstrated that NSCs, like their ectodermal progenitors, have low levels of surface glycosylation, a feature associated with increased susceptibility to SARS-CoV-2 infection relative to non-ectodermal cell types (Song et al., 2025). NSCs also utilize both entry pathways, with a marked preference for endocytosis during Omicron pseudoparticle exposure. Our data collectively demonstrate that both ectoderm and its NSC derivatives are highly susceptible to SARS-CoV-2 infection, in part due to low levels of glycosylation on their cell surfaces. This stresses the critical role of glycosylation in modulating viral susceptibility during early human development and highlights the need for further investigation into glycan-mediated host–virus interactions in embryonic and fetal contexts.

Consistent with our previous study (Song et al., 2025), enzymatic removal of the protective glycocalyx using neuraminidase significantly increased infection across all three types of spike pseudoparticles. This emphasizes the importance of surface glycosylation in modulating viral susceptibility. Other studies have further shown that glycosylation of viral spike protein and the ACE2 host receptor can influence infection. For example, glycan shielding on ACE2 may obscure key binding sites, thereby reducing viral entry efficiency (Yang and Rao, 2020; Zhao et al., 2020).

In our prior study, the presence of a robust glycocalyx on endodermal and mesodermal cells significantly diminished the efficacy of certain inhibitors (Song et al., 2025). For example, OcTMAB demonstrated robust efficacy with hESCs and endoderm; however, it was ineffective in mesoderm, where glycosylation was abundant. In mesoderm, the glycocalyx may have impeded inhibitor access to TMPRSS2 and endocytic machinery at the cell surface, thereby lowering its efficacy. Despite the high activity of TMPRSS2, the endoderm was exclusively infected via endocytosis (Song et al., 2025), perhaps because inhibitor penetration to the TMPRSS2 was impeded by the glycocalyx. The variable efficacy of the inhibitors may be related to their ability to reach the cell surface and affect TMPRSS2 and endocytosis. These observations highlight the importance of glycan architecture in governing both viral entry and the pharmacodynamics of entry-targeting therapeutics.

In summary, our findings support the conclusion that SARS-CoV-2 poses a significant risk to the developing nervous system in humans. Building on our prior work, this study demonstrates that the ectoderm and its NSC derivatives exhibit higher susceptibility to SARS-CoV-2 infection than hESCs, endoderm, and mesoderm. The susceptibility of the ectoderm and its derivatives is primarily attributed to the utilization of two viral entry pathways and the sparse glycocalyx on ectodermal and NSC cell surfaces, which facilitates viral access to the host receptors. The efficacy of entry inhibitors varied with filipin emerging as the most potent agent in NSCs and a potential candidate for further translational investigation. Given the potential for *in utero* exposure to disrupt neurodevelopment, longitudinal monitoring of offspring born to SARS-CoV-2–infected mothers is needed to identify and address cognitive and neurological sequelae.

## Data availability

No datasets were generated or analyzed during the current study.

## Competing interests

The authors declare no competing interests.

## Acknowledgements

We thank the UCR Stem Cell Core for providing access to the Novocyte Flow cytometer. Supported in part by a Committee on Research grant from the UCR Academic Senate, Academic Merit Fellowship, Yvonne Danielson Endowed Graduate Award, and the Dissertation Completion Fellowship Award from the UCR Graduate Division.

## Author contributions

A.S., C.Z., and P.T. contributed to data interpretation and manuscript writing. A.S. and P.T. were responsible for conception and manuscript editing of the submitted version. A.S. and C.Z. contributed to the sample preparation, data collection, and processing. P.T. was responsible for project administration and funding acquisition.

---

**Supplementary Table S1.** List of abbreviations and their definitions.

| Abbreviation | Definition |
| --- | --- |
| ACE2 | Angiotensin-Converting Enzyme 2 |
| ANOVA | Analysis of Variance |
| COVID-19 | Coronavirus Disease 2019 |
| DMEM | Dulbecco's Modified Eagle Medium |
| DMSO | Dimethyl Sulfoxide |
| DTT | Dithiothreitol |
| hESCs | human Embryonic Stem Cells |
| mβCD | methyl-β-cyclodextrin |
| MiTMAB | Myristyl trimethyl ammonium bromide |
| NSCs | Neural Stem Cells |
| OcTMAB | Octadecyltrimethylammonium Bromide |
| PBS | Phosphate-Buffered Saline |
| PFA | Paraformaldehyde |
| SARS-CoV-2 | Severe Acute Respiratory Syndrome Coronavirus<br>2 |
| TMPRSS2 | Transmembrane Protease, Serine 2 |

## Notes

### Competing Interest Statement

The authors have declared no competing interest.

